# Sequencing of individual barcoded cDNAs on Pacific Biosciences and Oxford Nanopore reveals platform-specific error patterns

**DOI:** 10.1101/2022.01.17.476636

**Authors:** Alla Mikheenko, Andrey D Prjibelski, Anoushka Joglekar, Hagen U Tilgner

**Author notes:** Contributed equally (Alla Mikheenko, Andrey D Prjibelski).

## Abstract

Long-read transcriptomics requires understanding error sources inherent to technologies. Current approaches cannot compare methods for an individual RNA molecule. Here, we present a novel platform comparison method that combined barcoding strategies and long-read sequencing to sequence cDNA copies representing an individual RNA molecule on both Pacific Biosciences and Oxford Nanopore. We compared these long reads pairs in terms of sequence content and splicing structure. Although individual read pairs show high similarity, we found differences in (i) aligned length, (ii) TSS and (iii) polyA-site assignment, and (iv) exon-intron structures. Overall 25% of read pairs disagreed on either TSS, polyA-site, or a splice site. Intron-chain disagreement typically arises from alignment errors of microexons and complicated splice sites. Our single-molecule technology comparison revealed that inconsistencies are often caused by sequencing-error induced inaccurate ONT alignments, especially to downstream GTNNGT donor motifs. However, annotation-disagreeing upstream shifts in NAGNAG acceptors in ONT are often confirmed by PacBio and thus likely real. In both barcoded and non-barcoded ONT reads, we found that intron number and proximity of other GT/AGs better predict inconsistency with the annotation than read quality alone. We summarized these findings in an annotation-based algorithm for spliced alignment correction that improves subsequent transcript construction with ONT reads.

## Introduction

Long-read sequencing is increasingly used in transcriptomics, including its application to barcoded unique molecules (Gupta et al. 2018; Singh et al. 2019), giving single-cell and spatially resolved long-read transcriptomes. Various technologies currently exist, including Pacific Biosciences (Eid et al. 2009; Koren et al. 2012; Sharon et al. 2013; Au et al. 2013; Tilgner et al. 2014; Weirather et al. 2015), Oxford Nanopore(Oikonomopoulos et al. 2016; Byrne et al. 2017), as well as linked-read technologies. Linked-read technologies for RNA were originally represented either by synthetic long reads (SLRs) (Tilgner et al. 2015) or usually more sparsely covered 10x Genomics linked-reads (Tilgner et al. 2018), although more recently other linked-read technologies have emerged (Wu et al. 2019; Chen et al. 2020). Furthermore, for all these platforms a variety of protocols either exists or can be imagined. Comparing the accuracy of these distinct approaches is therefore fundamental in modern transcriptomics just as it has been fundamental for short-read sequence analysis (Engström et al. 2013; Steijger et al. 2013; Li et al. 2014b, 2014a).

A drawback of commonly used strategies is their lack of single-molecule resolution. For example, percent-spliced-in (PSI)-values (Wang et al. 2008) of splices sites or transcript-per-million (TPM) (Wagner et al. 2012) values can easily be compared between multiple strategies. However, these approaches do not allow to compare the accuracy of different strategies for a single molecule. At the same time, recent advances in long-read sequencing of barcoded molecules allow the identification of the exact molecule that is being sequenced. This information can be used to compare how different technologies behave for the same molecule.

Here, we compare cDNAs that are barcoded by their single-cell (ScISOr-Seq) of the origin or their spatial location (Sl-ISO-Seq), sequenced on both the Pacific Biosciences Sequel II (circular consensus reads, “PacBio” from here on) and Oxford Nanopore Promethion (“ONT”) systems (Joglekar et al. 2021). Thus, PCR copies of the same reverse-transcription event are sequenced on both platforms (“RT read pairs”), which enables their comparison for individual RNA molecules. A reverse-transcription event is identified by the combination of (i) single-cell/spatial location barcode, (ii) a unique molecular identifier (UMI), and (iii) the gene the molecule is mapped to (Fig. 1a). In order to have the highest confidence barcodes and ONT-PacBio read pair correspondences, we only use perfect matching for barcode and UMI detection. Thus, we compared sequencing errors and intron structure in identified RT read pairs. In brief, we demonstrate that PacBio reads are significantly more accurate and typically capture slightly longer transcript portions than ONT reads. While ONT and PacBio reads from RT pairs often agree on splicing structure, inconsistencies mostly arise from inexact ONT alignments. We also show that observations made for RT read pairs are generally true for non-barcoded reads and can be exploited to perform correction of Nanopore spliced alignments.

**Fig. 1.**
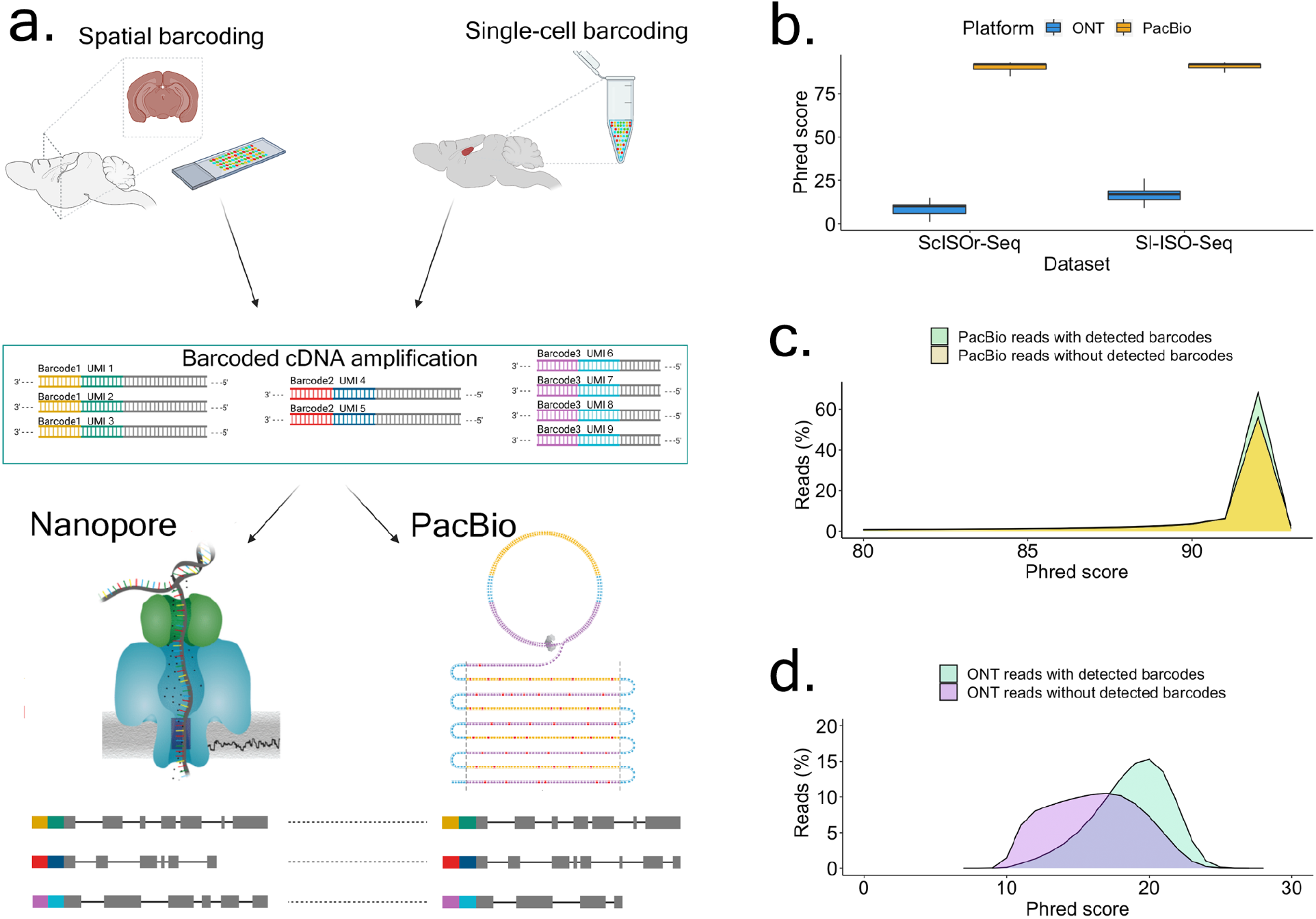
Outline and primary read characteristics. **a.** Individual reverse transcription events turn an individual RNA molecule into a barcoded RNA-cDNA hybrid, which is amplified into many cDNA molecules that carry the same barcode and UMI. We carried this process out in two distinct ways - by single-cell 10xGenomics barcoding as well as by spatial 10xGenomics Visium barcoding. Aliquots of these cDNAs are then sequenced on PacBio and ONT. Using the identity of barcode and UMI, we can detect individual RNA molecules, whose cDNA copies have been sequenced on both ONT and PacBio. We refer to these read pairs as RT read pairs. **b.** Comparison between Phred scores of PacBio CCS and ONT reads from both datasets. **c.** Phred score distribution for PacBio CCS reads from Sl-ISO-Seq dataset with (light green) and without (yellow) detected barcodes. **d.** Phred score distribution for Oxford Nanopore reads from Sl-ISO-Seq dataset with (light blue) and without (purple) detected barcodes.

## Results

### Molecule identification discards lower quality ONT reads

We first compared Phred scores of all individual reads sequenced on both platforms independently of molecule detection. The Phred quality score of a base indicates the probability of a base being called correctly. The Phred quality score of a read is computed as the average across Phred scores of all read bases. Unsurprisingly, PacBio showed dramatically higher Phred scores than ONT both in single-cell and in spatial data (Fig. 1b). Importantly, since we conservatively analyzed only perfectly matched barcodes, ONT reads with barcodes had significantly higher Phred scores than those without (Fig. 1d, two-sided Wilcoxon-rank-sum p-value=2.4*10^-9^), while no significant difference was detected for PacBio (Fig. 1c, two-sided Wilcoxon-rank-sum p-value=0.2). Similar observations were made on the single-cell dataset, although the difference for ONT reads was even more dramatic (Fig. S1a-b). Overall, barcode detection (Supplementary Note “Experimental details”) in spatial data yielded 2,873,455 PacBio reads (of 3,371,331 - 85%) and 12,153,599 ONT reads (of 73,181,790 - 16%). Although ONT essentially has a significantly deeper sequencing depth than PacBio, the difference between the number of sequenced reads is not so dramatic considering only reads with detected barcodes. If mismatches in barcodes were allowed, it is likely that more Nanopore reads could be recovered, although at the risk of introducing more false positives.

### Sequence comparison using reference-based and reference-free alignments

Our further analysis mostly relies on read mappings, so at first we aligned reads using different tools: widely popular minimap2 (Li 2018) and specialized transcriptome aligners deSALT (Liu et al. 2019) and GraphMap2 (Marić et al. 2019) to exclude a possible bias. Although all three aligners yield largely similar results, GraphMap2 produced slightly shorter alignments than minimap2 and deSALT with more prominent differences between aligned lengths of PacBio and ONT reads. In addition, GraphMap2 produced the least number of RT read pairs because a PacBio read and an ONT read sharing the same barcode and UMI were more frequently mapped to the different genes (Supplementary Table 3).

Below we also show that alignments generated by deSALT and especially Graphmap2 show more discordance between PacBio and ONT reads in terms of splice sites than alignments produced by minimap2 (Fig. S6) and minimap2 has significantly better performance in terms of isoform detection (Section “Splice site correction improves transcript discovery precision”, Table 1). Of note, STARlong (Dobin et al. 2013) has strong performance for PacBio reads, but is not optimised for error-prone ONT data. To enforce the same strategy for PacBio and ONT, here we use minimap2 for alignment.

**Table 1.** Precision, sensitivity, and false positive rates of StringTie2 results on the simulated ONT dataset aligned with different strategies. The best values across different methods are highlighted in bold.

|  | All isoforms |  | Known isoforms |  | Novel isoforms |  |  |
| --- | --- | --- | --- | --- | --- | --- | --- |
|  | Sensitivity | Precision | Sensitivity | Precision | Sensitivity | Precision | # false transcripts |
| deSALT without annotation | 36.5 | 24.1 | 39.1 | 80.9 | 17.0 | 2.3 | 38,209 |
| minimap2 without annotation | 84.2 | 58.0 | 86.5 | 88.0 | 57.7 | 14.1 | 18,683 |
| minimap2 with annotation | 86.4 | 67.7 | 88.0 | 89.4 | **64.2** | 21.9 | 12,155 |
| minimap2 with annotation and additional correction | **89.8** | **73.6** | **92.7** | 88.7 | 64.0 | **29.1** | **8,281** |

Mapping RT read pairs (total 54,752 in Sl-ISO-Seq dataset, 274,287 in ScISOr-Seq) to the genome with minimap2 (Li 2018) showed highly correlated alignment lengths (Figs. 2a and S2a). PacBio reads were significantly longer than their ONT counterparts in ScISOr-Seq and, although less pronounced, in Sl-ISO-Seq data, but differences were small compared to the entire read (Fig. 2b). However, in terms of absolute numbers, the difference is notable: 70% of read pairs have a longer aligned portion (median: 14bp) in the PacBio and 27% in the ONT read (median 8bp, Fig. 2c). In summary, in comparison to read length, the differences are minor. However, 8-14bp sequences can harbor important elements such as polyA signals, protein/miRNA-binding sites, microexons, or G-quadruplexes (Lee et al. 2020).

**Fig. 2.**
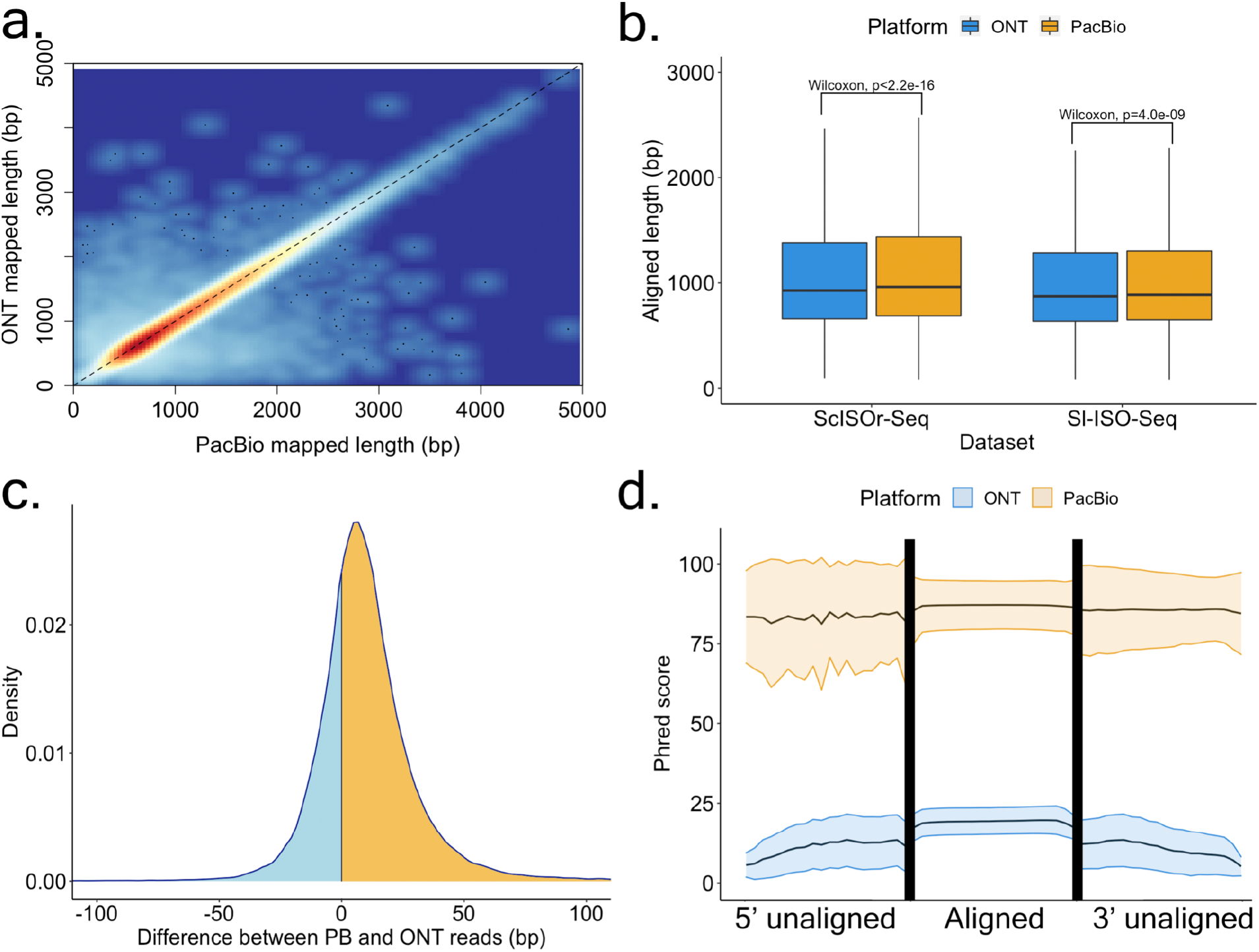
Alignment characteristics of RT read pairs. **a.** Heatscatter plot showing aligned lengths of respective PacBio read (X-axis) and ONT read (Y-axis) from the RT read pair after mapping to the genome using minimap2 (Sl-ISO-Seq dataset, Spearman Rho: 0.96, p<2.2*10^-16^). **b.** Comparison between aligned lengths of PacBio and ONT reads for both datasets after mapping to the genome using minimap2. **c.** Density plot showing the difference between aligned lengths of PacBio read and ONT read from the RT read pair after mapping to the genome using minimap2 (Sl-ISO-Seq dataset). Cases when PacBio alignment is longer correspond to the yellow area under the curve, otherwise represented by the blue area. **d.** Mean Phred score distribution along the read for aligned (middle) and unaligned (left and right) parts of PacBio (yellow) and ONT (blue) reads based on a (reference-free) pairwise Smith-Waterman alignment of the PacBio and Nanopore reads from the RT read pair (Sl-ISO-Seq dataset). Lower and upper bounds represent the standard deviation of the Phred score distribution.

To delineate common and diverging sequences in reads from RT pairs, we aligned them using the Smith-Waterman algorithm to each other. Of note, unlike modern mapping algorithms based on heuristics, Smith-Waterman is an exact solution and not subject to future improvements. We divided each alignment pair into an unaligned 5’ part, an aligned portion, and an unaligned 3’ part. In the spatial data, in all three areas, PacBio showed much higher read-wise Phred scores than ONT and PacBio qualities slightly drop as it enters the non-aligned 5’ area, while for single-cell data PacBio qualities in the 5’ unaligned portion remained constant and comparable to the aligned area. In both datasets, ONT qualities deteriorate in the 5’ unaligned area and gradually decrease in the unaligned 3’ area (Figs. 2d and S2b).

### Sequencing error rates and k-mer identity in alignments to the genome

Deducing sequencing errors from alignments is difficult because, in addition to sequencing and alignment errors, PCR errors, SNVs and mutations cause a divergence between reads and the genome. We used a three-way comparison of paired PacBio and ONT reads as well as the genome to define a “ground truth” using a majority call among all three sources and delineate error patterns as a divergence from this ground truth (Methods). Unsurprisingly, ONT has more errors than PacBio (Fig. S3a). For PacBio, deletions or mismatches are ~3-fold less abundant than insertions, whose frequency increases towards read ends (Fig. S3b). ONT behaves very differently: deletions dominate over insertions and mismatches, and all error types decrease towards alignment ends (Fig. S3c). 41% of PacBio errors and only 23% of ONT errors occur in homopolymers (Fig. S3d). For PacBio, indels within homopolymers are more prevalent than mismatches, and slightly more errors occur in homopolymers towards the 3’end (Fig. S3e). For ONT, similar trends were observed, although the insertions are less biased towards homopolymers than in PacBio (Fig. S3f). In summary, PacBio has fewer errors than ONT, but a large fraction of PacBio errors occur in homopolymers, while ONT errors mostly arise from other areas. Similar observations were obtained for the older ScISOr-Seq dataset, although the overall error rate in ONT reads was higher (Fig. S4a-d).

Alignments to a genome often employ seeding through matching k-mers. We considered the 14-mer identity, a commonly used k-mer for example in minimap2 (Li 2018), and analyzed each exon alignment separately. As expected, 14-mer identity was lower than single-base identity and specifically affected ONT reads (Fig. S3g). Unexpectedly, for ONT data, we found lower 14-mer identities for slightly longer exons (two-sided Wilcoxon-rank-sum test p=0.005, 1-20bp vs. 21-50bp exons) and a tighter distribution. A possible explanation is that short exons may become unmappable with few sequencing errors. For PacBio, a more intuitive pattern emerged: longer exons have a higher 14-mer identity than shorter exons (Fig. S3g). After homopolymer compression (Methods), 14-mer identity reached almost 100% for PacBio regardless of exon-length. However, for ONT, compression causes high variability in short exons (<21bp): while the median increased to 100%, the 1st quartile decreased to 0%, as a single error in a short exon can affect all 14-mers (Fig. S3h). Broadly similar observations were made for Sc-ISOr-Seq data (Fig. S4e-f). Thus, homopolymer compression should be applied to ONT reads with care.

### Three-way comparison of annotation, ONT and PacBio shows differences at exon and splice-site calling

Long-read experiments regularly uncover many isoforms that are inconsistent with annotations (Sharon et al. 2013; Au et al. 2013; Tilgner et al. 2014, 2015; Oikonomopoulos et al. 2016; Tardaguila et al. 2018; Tung et al. 2019; Kovaka et al. 2019; Tang et al. 2020; Wyman et al. 2020). While for short-read experiments, splice site identification and the following splice site quantification has been addressed with large success (Vaquero-Garcia et al. 2016), long-read based annotation-inconsistent isoforms could be truly novel or simply false. This question can only be conclusively answered for a single molecule as a correct alignment for one molecule can be false for another molecule. Here, we exploit RT read pairs to evaluate inconsistencies between alignments and annotation - and if they are observed on both platforms.

We considered 22,600 and 48,993 RT read pairs, where at least one of the PacBio/ONT reads is assigned to a known TSS (Lizio et al. 2015) and polyA site (Herrmann et al. 2019) respectively (Methods). Indeed, all barcoded reads have a polyA tail, which creates a bias towards 3’completeness in RT read pairs, and, thus, a known polyA site is assigned for a significantly larger portion of reads than a TSS. Moreover, in 95% of RT pairs both PacBio and ONT are assigned to the same annotated polyA site, while agreement on TSS is lower (87%, Fig. 3a,b). For TSS assignment a significant portion of disagreeing pairs arises from unassigned ONT reads (8% of all RT pairs), which suggests that 5’ truncation of PacBio reads is less frequent. Unexpectedly, we noticed that the results differ for spliced and unspliced reads. While spliced reads are assigned to TSS more often, the percentage of RT pairs with both reads spliced and assigned to different TSS is also higher (3.2% for spliced reads vs 1.7% for unspliced). This may potentially be explained by short starting exons misaligned in ONT reads. For polyA sites, however, the difference between spliced and unspliced reads is marginal (Fig. 3c).

**Fig. 3.**
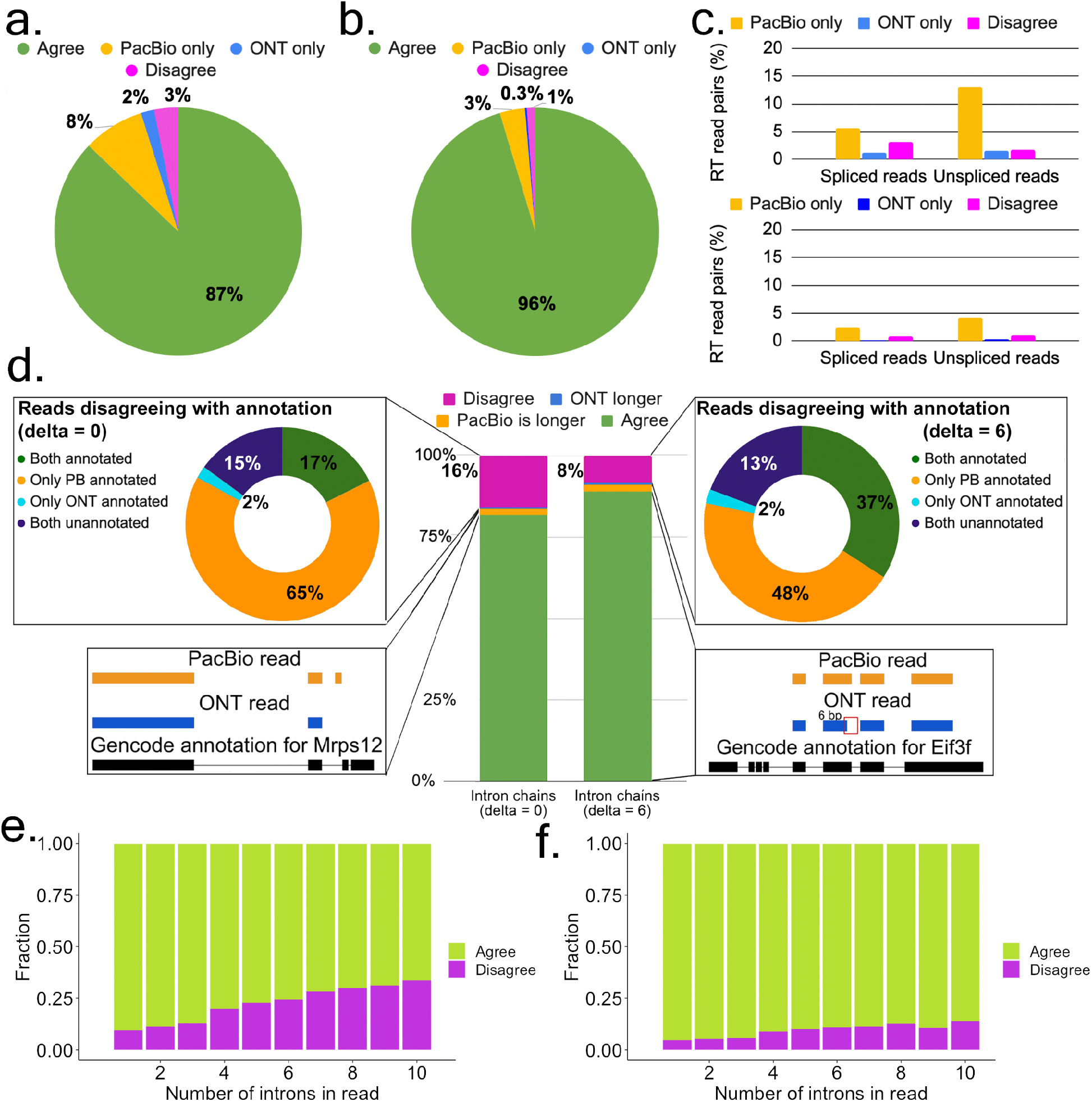
Agreement in RT read pairs of Sl-ISO-Seq data. **a.** Fractions of TSS assignments that agree (green), disagree (magenta), or found only in one read (blue for ONT, yellow for PacBio) from the RT pair. **b.** Same as Fig. 3a, but for Poly-A sites. **c.** Percentage of RT read pairs that disagree on the assigned TSS (top) and Poly-A site (bottom): only PacBio read was assigned (yellow), only ONT (blue), both assigned but to different sites (magenta). **d.** *Middle:* percentage of RT read pairs that agree (green), disagree (magenta), or one chain being longer (blue for ONT, yellow for PacBio) when splice junctions are compared precisely (left, delta=0bp) or inexactly (right, delta=6bp). *Top left:* classification of disagreeing intron chains from RT read pairs with respect to the reference annotation (delta=0bp): both are inconsistent with the annotation (dark blue), both correspond to known (different) transcripts despite the disagreement (green), PacBio is consistent with the annotation while ONT is not (yellow) and vise versa (light blue). *Top right:* classification of disagreeing intron chains with respect to the reference annotation using inexact comparison (delta=6bp). *Bottom left:* an example of agreeing intron chains from an RT read pair, in which PacBio intron chain is longer (Mrps12 gene). *Bottom right:* An example of intron chains from an RT read pair that have a 6bp difference in the donor site of the second intron. Comparing intron chains with delta=6bp classifies them as agreeing (Eif3f gene). **e.** Fraction of agreeing (green) and disagreeing (magenta) intron chains with respect to intron chain length when compared precisely (delta=0bp). **f.** Same as Fig. 3e, but with delta=6bp.

We then considered for each read mapping the list of all its introns, which we refer to as an intron chain. In 81.8% of RT read pairs (of 28,330 total pairs in which both reads are spliced), the PacBio and the ONT read had identical intron chains. In 2% of cases, intron chains were not contradictory, but the PacBio read had extra intron(s) at its extremities, while ONT reads have longer intron chains only in 0.6% of pairs. In the remaining 15.7%, PacBio and ONT intron chains disagree with each other. When compared to the annotation, in 17% of the disagreeing RT read pairs both intron chains corresponded to different known transcripts (green in Fig. 3d, top left), while in 15% of cases both were inconsistent with the annotation (dark blue in Fig. 3d, top left). In these cases, it is not easy to ascertain which mapping is true. However, in a large fraction (65%) of the disagreeing pairs, the PacBio mappings were consistent with the annotation, while ONT mappings were not (yellow in Fig. 3d, top left). In this case, assuming that the PacBio intron chain is in fact correct appears more parsimonious than the contrary.

The above statements are based on a single-base interpretation of PacBio and ONT splice sites (delta=0bp, Methods). To account for slight shifts in splice site mapping, we explored inexact intron chain comparison, in which junctions are considered equal if the distance between them does not exceed 6bp (delta=6bp). This reduced disagreements between paired PacBio and ONT reads by 48%. Among the remaining disagreements, in nearly half of the cases the PacBio mapping corresponded to an annotated transcript, while the ONT read did not (Fig. 3d middle and right). Overall, 46% of ONT and 15% of PacBio mappings inconsistent with the annotation at delta=0bp were reclassified as annotated with delta=6bp.

We hypothesized that the fraction of disagreeing RT read pairs would increase with the number of splice sites per read. Indeed, reads with ≥8 introns disagreed with its pair ~3-fold more often than reads with 2 introns. However, even read pairs with ≥8 introns still agreed in 70% of cases (Fig. 3e). Using delta=6bp reduces disagreements but roughly preserves the trend (Fig. 3f). These observations suggest that other factors, beyond the intron chain length, also influence disagreements between PacBio and ONT reads. We, therefore, investigated sequence characteristics of disagreeing introns.

### Sequence characteristics underlying disagreeing within RT read pairs

We then analyzed alternative splicing events in 23,356 pairs of PacBio and ONT reads where both reads in a pair were spliced, uniquely mapped, and unambiguously assigned to an isoform (Methods). Exon skipping with respect to the identified isoform was observed 461 times (1.9%) in ONT data and only 45 times (0.2%) in PacBio data (Fig. 4a). Short exons were skipped more often, although much less frequently for PacBio than for ONT. ONT exon skipping typically affects exons ≤40bp, however PacBio rarely skips exons longer than 15bp (Fig. 4b). Minimap2 employs exact matching of certain k-mers with default k=15 (Li 2018) followed by dynamic programming. However, sequencing errors, especially in ONT data, may cause the above short exons to be missed.

**Fig. 4.**
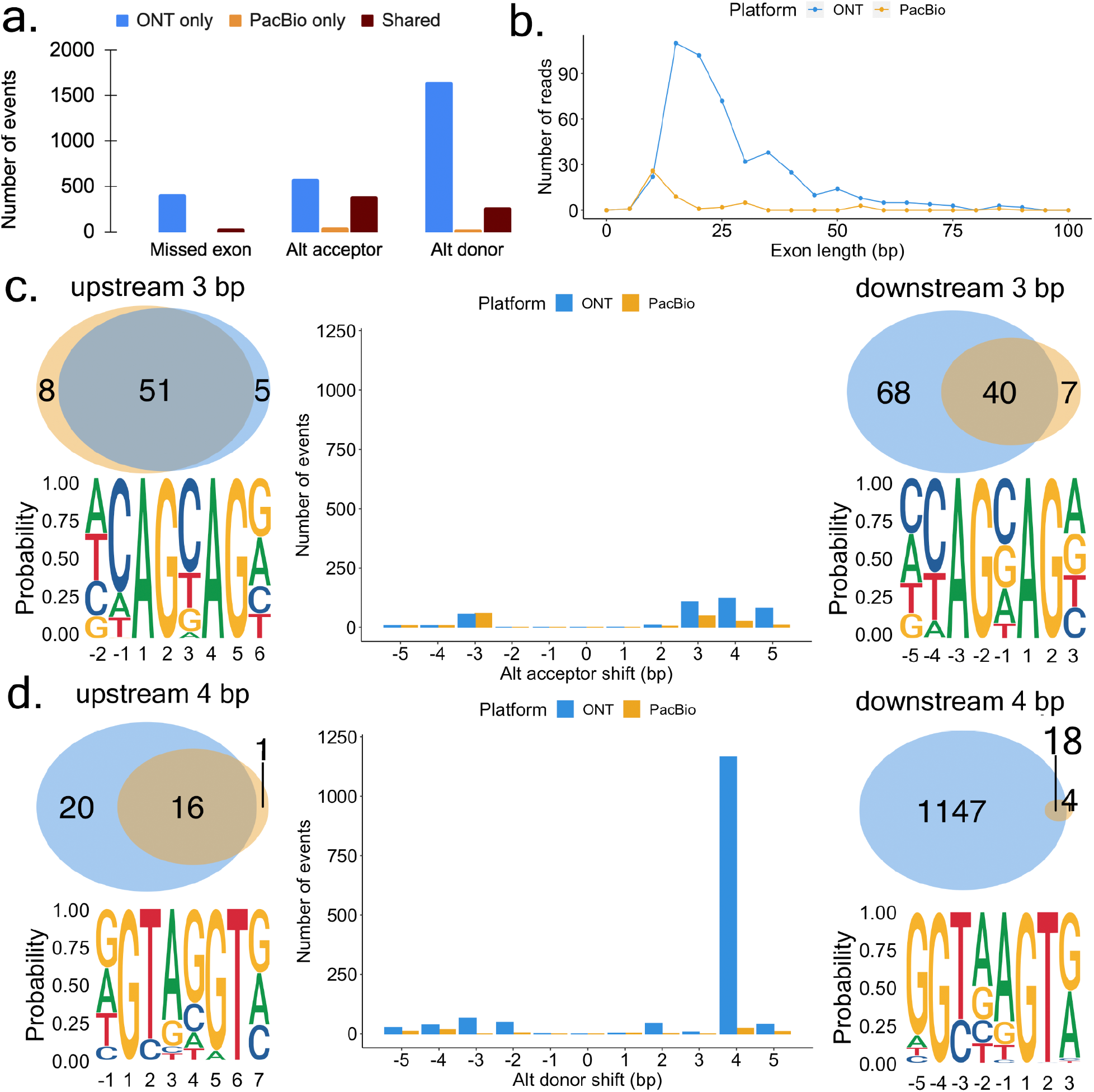
Exon and splice site characteristics underlying disagreements between PacBio and Nanopore in Sl-ISO-Seq data. **a.** Number of missed exons (left), alternative acceptors (middle), and donors (right) with respect to the reference annotation that occur only in ONT read (blue), only in PacBio read (yellow), and in both reads from RT pair (brown). **b.** Length distribution for skipped exons in PacBio reads (yellow) and ONT reads (blue). **c.** *Middle:* number of alternative acceptor sites in PacBio reads (yellow) and ONT reads (blue) with respect to distance from the annotated acceptor site. *Top left:* Venn diagram for 3bp upstream alternative acceptor sites in PacBio (yellow) and ONT reads (blue) from RT pair. *Bottom left:* Nucleotide frequency for loci where 3bp upstream acceptor sites occur. *Top right:* Venn diagram for 3bp downstream alternative acceptor sites in PacBio (yellow) and ONT reads (blue) from RT pair. *Bottom right:* Nucleotide frequency for loci where 3bp downstream acceptor sites occur. **d.** *Middle:* number of alternative donor sites in PacBio reads (yellow) and ONT reads (blue) with respect to distance from the annotated donor site. *Top left:* Venn diagram for 4bp upstream alternative donor sites in PacBio (yellow) and ONT reads (blue) from RT event pair. *Bottom left:* Nucleotide frequency for loci where 4bp upstream donor sites occur. *Top right:* Venn diagram for 4bp downstream alternative donor sites in PacBio (yellow) and ONT reads (blue) from RT pair. *Bottom right:* Nucleotide frequency for loci where 4bp downstream donor sites occur.

Alternative acceptors (4.2% of ONT reads; 2% of PacBio reads), and, comparatively more often, alternative donors (8.3% for ONT and 1.3% for PacBio) were more common than exon skipping (Fig. 4a). Thus, overall 11.6% of ONT reads and 3.7% of PacBio reads showed one or more discrepancies to the annotation. We found similar trends in the single-cell data, albeit with higher inconsistencies for ONT data (Fig. S7a-b). Discrepancies between an alignment and annotation found in both PacBio and ONT likely are novel isoforms. Such cases generally exceed discrepancies supported by PacBio only (Fig. 4a).

From here on, we only consider canonical introns (GT-AG). We found that inconsistent acceptors were usually shifted by 3-5bp downstream in ONT only, while 80% upstream 3bp-shift (“NAGNAG”) acceptors were supported by both PacBio and ONT. Thus, downstream ONT acceptor shifts are questionable, while upstream NAGNAG shifts appear often true (Fig. 4c). For inconsistent donors, a downstream shift of 4bp predominantly occurred for ONT data and with a much smaller overlap between both technologies than for the acceptor sites (Fig. 4d). Such shifts are caused by misalignment at the commonly known GTNNGT donor motif (Wang and Ruvinsky 2010). In summary, downstream 4bp shifts from an annotated donor observed in ONT are doubtful, while 3bp shifts from an annotated acceptor harbor a significant number of trustworthy novel splice sites. Broadly similar observations were made with Sc-ISOr-Seq data (Fig. S7c-d).

It is worth noting that modern transcriptome aligners can use annotated splice junctions. This dramatically reduced the discrepancies between ONT and the annotation but had a marginal effect on PacBio and cases, for which ONT and PacBio agreed. Importantly, using the annotation causes PacBio to have more alignment inconsistencies than ONT, possibly because of overcorrection of ONT splice sites to the annotation (Fig. S7e-h).

### Extrapolating characteristics observed in barcoded read pairs to non-barcoded ONT reads

The observations described above are based on RT event read pairs, which require a detected barcode and UMI in both the PacBio and the ONT read. However, as opposed to PacBio data, ONT reads with barcodes have higher Phred scores than those without (Fig. 1d). To understand quality effects on read characteristics, we analyzed the entire Sl-ISO-seq ONT data, which mimics non-barcoded transcriptomic experiments. We observed that the aligned length increased from 532bp for read-wise Phred-score=10 reads to 815bp for Phred-score=20 (Fig. 5a). Similarly, high-quality reads had one more detected intron on average: 2.7 and 3.7 introns for spliced reads with Phred score 10 and 20 respectively (Fig. 5b).

**Fig. 5.**
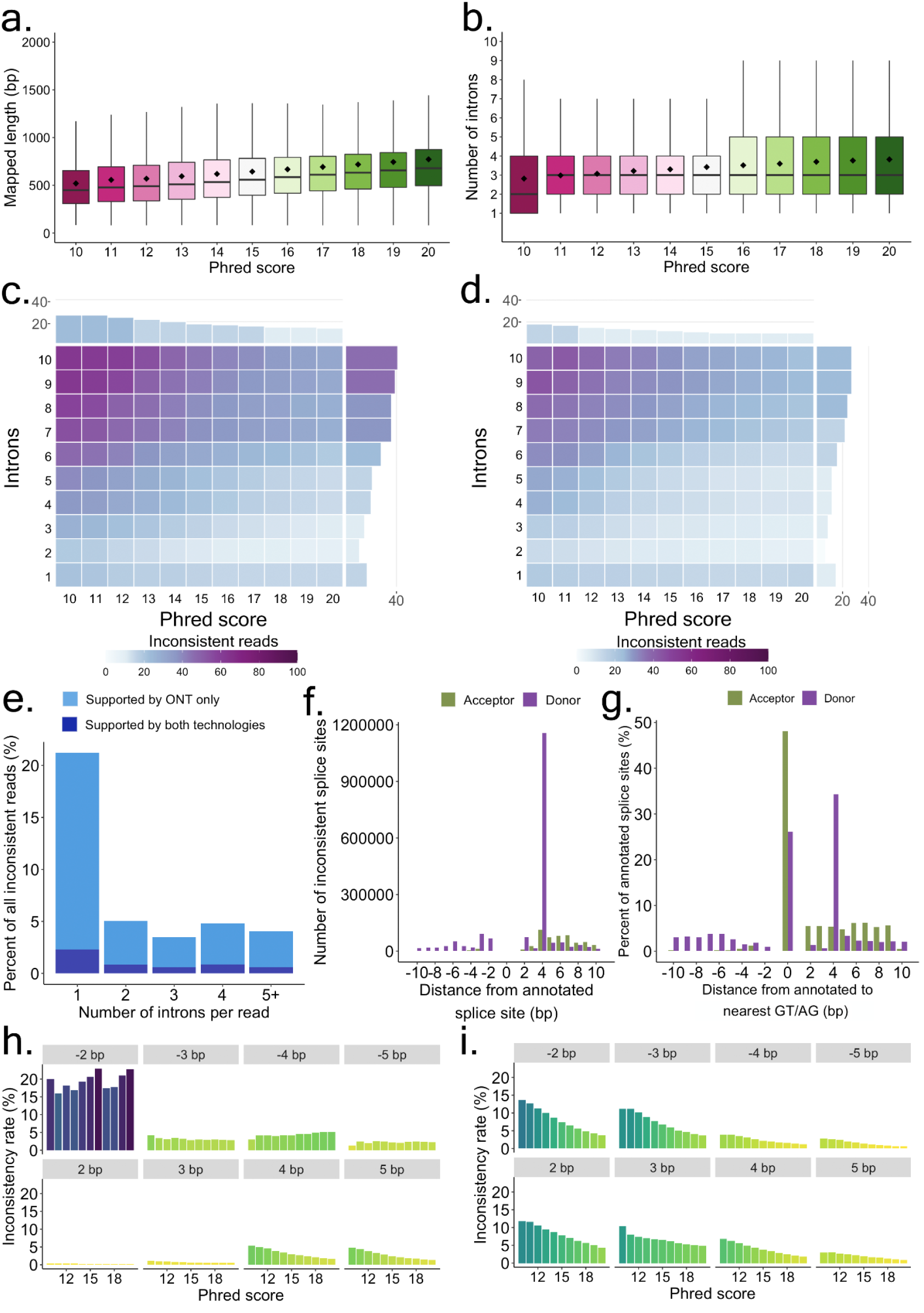
Characteristics of all including non-barcoded ONT reads for Sl-ISO-Seq data. **a.** Aligned read length with respect to read Phred score (average across all read bases) for all ONT reads from Sl-ISO-Seq dataset. **b.** Read intron chain length with respect to read Phred score for all ONT reads from Sl-ISO-Seq dataset. **c.** Heatmap showing average inconsistency rate between read intron chains and annotated intron chains (exact comparison, delta=0bp) with respect to read Phred score (X-axis) and intron chain length (Y-axis) for all ONT reads from Sl-ISO-Seq dataset. Barplot at the top (on the right side) summarizes the inconsistency rate with respect to only the Phred score (only intron chain length). Purple corresponds to a higher inconsistency rate, while light blue indicates a lower inconsistency. **d.** Same as Fig. 5c, but using inexact intron chain comparison (delta=6bp). **e.** Histogram showing a fraction of inconsistent ONT reads that have at least one intron entirely contained inside an annotated exon. Dark blue represents reads for which the contained intron is supported by at least one PacBio read, while light blue corresponds to the rest of ONT reads. **f.** Number of inconsistent donor (purple) and acceptor (green) splice sites in ONT reads from Sl-ISO-Seq dataset with respect to the distance from the annotated splice site. **g.** Percentage of annotated canonical donor (green) and acceptor (purple) splice sites with respect to distance to the nearest canonical dinucleotides (GT for donors, AG for acceptors). 0 corresponds to the case when no canonical dinucleotides were detected in the vicinity of 10bp. **h.** Inconsistency rates of individual acceptor splice sites in ONT reads from Sl-ISO-Seq dataset with respect to read Phred score. Each histogram represents splice sites with a certain distance from the annotated splice site (grey bars on top). **i.** Same as Fig. 5h, but for donor splice sites.

Furthermore, we compared each read intron chain against annotated transcripts (Methods). The inconsistency rate goes down with higher Phred score: 28% of reads are inconsistent with the annotation for Phred-score=10 reads (*n*=206,675), but only 15% are inconsistent for Phred-score=20 reads (*n*=2,667,759, Fig. 5c). Moreover, as found previously (Tilgner et al. 2013; Sharon et al. 2013; Tilgner et al. 2015; Au et al. 2013), inconsistency increases with longer intron chains. I.e., 40% of reads with ≥7 introns are inconsistent, ~2.7-fold more than for reads containing ≤3 introns (~15%). Similar trends were observed when intron chains were compared using delta=6bp, although overall inconsistency rates dropped (Fig. 5d). Interestingly, the lowest inconsistency rate is observed for reads with 2 introns (11%), rather than for single-intron reads (17%). While this observation was previously reported, no explanation was found (Tilgner et al. 2013; Sharon et al. 2013; Tilgner et al. 2015). We hypothesized that aligners can arbitrarily split a read into two exons, while such splits into 3 or more exons are less likely. Additional analysis (Methods) showed that 19% of inconsistent single-intron alignments have their intron entirely within an annotated exon, ~5-fold more than for alignments with 2 introns (3.5%, Fig. 5e). Moreover, only 12% of such mappings are supported by PacBio reads. Thus, some inconsistent single-intron mappings likely occur due to misalignments, rather than represent real alternative isoforms.

We further examined individual splice sites by considering all canonical introns in ONT alignments that matched annotated canonical introns with a loose threshold of 10bp. Similar to barcoded ONT reads, mapped acceptors are rarely shifted upstream and are more consistent with the annotation than donors, which have a dominant 4bp downstream shift caused by the GTNNGT motif (Fig. 5f). To understand the major source of these shifts, we computed distances from all annotated splice sites to the nearest AG/GT (Fig. 5g). As these distances strongly resemble the distribution of ONT shifts, we hypothesized that shifts may depend on the proximity of AG/GT in general, rather than a particular motif.

Thus, we analyzed the inconsistency rate of annotated splice sites with respect to (i) the nearest GT/AG and (ii) read quality (Methods). Donor inconsistency strongly depends on both read quality and distance to the nearest GT (Fig. 6i). For example, the probability of a downstream 4bp shift caused by GTNNGT motif is ~2 times lower than of a downstream 2bp shift near GTGT (3.06% and 6.48% respectively), despite GTNNGT being vastly more frequent than GTGT overall (36,663,585 and 1,066,886 of total detected splice sites near respective motif). Acceptor sites, on the other hand, show visible quality dependency only for downstream shifts, most likely due to rare occurrences of upstream AGs. The upstream shifts are dominated by NAGNAG acceptors, which show only a 1.5-fold decrease in inconsistency rate between reads with Phred-score 10 and 20 (4.25% vs. 2.86%). As this difference is noticeably smaller than that for all acceptors (~4.4-fold, 3.13% vs 0.71%), in conjunction with our RT read pair analysis, it suggests that a portion of the upstream NAGNAG shifts may be real.

In summary, due to the elevated number of sequencing errors, ONT read alignments obtained with minimap2 may not provide exact splice site coordinates, especially for the cases when (i) read has low quality, (ii) read spans multiple introns, and (iii) canonical dinucleotides are located nearby splice sites. To avoid potential misinterpretation of reads disagreeing with the annotated transcripts, one may treat such alignments with additional care or use inexact intron comparison when matching the annotation.

### Splice site correction improves transcript discovery precision

Based on the observations made for ONT data we implemented an algorithm for correcting splice junctions in individual reads with the aid of the annotation. This algorithm works with aligned reads and is capable of restoring (i) skipped short exons and (ii) incorrectly detected splice sites (Methods). To evaluate how the designed algorithm affects transcript discovery we simulated Nanopore reads with NanoSim (Hafezqorani et al. 2020), since for the real data the ground truth is unknown. Although one could use PacBio reads from the same RT event read pair for verification, the fraction of ONT reads having an RT pair is comparably low and is not suitable for transcript model construction.

Transcript discovery was performed using StringTie2 (Kovaka et al. 2019). To mimic real-life situations we removed some transcripts from the annotation before running our correction algorithm and StringTie2. The generated transcripts were matched against the set of “expressed” transcripts (known) and ones that were removed from the annotation (novel) using gffcompare (Pertea and Pertea 2020) to compute precision and recall of different approaches.

To understand the effect of splice site correction we generated read alignments using: (i) deSALT with default options; (ii) minimap2 with default options and without correction; (iii) minimap2 with the annotation and without correction; (iv) minimap2 with the annotation and the additional correction. Table 1 demonstrates that transcripts generated by StingTie2 using deSALT alignments have substantially lower quality compared to minimap2. At the same time, running minimap2 with the annotation greatly improves results for novel transcripts. However, additional correction with our algorithm makes further substantial improvement: overall sensitivity (precision respectively) increases by 3.4% (5.9% resp.) while the precision of novel transcripts improves by ~30% (from 21.9% to 29.1%) as the number of false positives among novel transcripts decreases 1.5 fold (from 12,155 down to 8,281).

Although both minimap2 and our correction algorithm used only information about known transcripts, this information aids the more precise detection of novel isoforms since they often share one or more exons with known isoforms.

## Discussion

Different long-read RNA approaches are increasingly used for isoform analysis (Koren et al. 2012; Au et al. 2013; Sharon et al. 2013; Tilgner et al. 2014, 2015; Oikonomopoulos et al. 2016; Tilgner et al. 2018; Garalde et al. 2018; Gupta et al. 2018; Depledge et al. 2019; Wang et al. 2019; Tardaguila et al. 2018; Volden et al. 2018; Tang et al. 2020; Sun et al. 2021; Joglekar et al. 2021; Hardwick et al. 2021). Therefore, understanding how each approach fares in detecting RNA traits is fundamental. Previous and current comparisons (Li et al. 2014b, 2014a; Weirather et al. 2017; Cui et al. 2020; Pardo-Palacios et al. 2021) of long-read technologies, while incredibly helpful due to their timeliness, size, and broad scope, could not compare multiple technologies for an individual RNA molecule. Here, we employed single-molecule barcoding technologies (Gupta et al. 2018; Joglekar et al. 2021) to sequence cDNA copies of single reverse transcription events on PacBio and ONT. Using perfectly matching barcodes and UMIs, we established the correspondence of a pair of Nanopore and PacBio reads to an individual RNA molecule. This procedure is highly specific while discarding doubtful PacBio-ONT read pairs, but causes a selection for higher-quality reads in ONT but not in PacBio (see Fig1c,d). We found important differences that can avoid misinterpretation of data and guide researchers in their choice of technology.

PacBio and ONT reads from an RT pair frequently differ in length, usually with up to 50 extra nucleotides in the PacBio read, which is small compared to the entire read. However, extra nucleotides in ONT also exist, although less frequently and fewer. These differences are small but can harbor polyA signals, Kozak sequences, or splicing factor binding sites. Additionally, we observed that PacBio extended more often to a known TSS and polyA site than ONT - a criterion important to defining complete isoforms. Importantly, despite an overall low error rate, a significant fraction of PacBio errors arises from homopolymers (up to 40%), while ONT shows more errors but with less bias towards homopolymers.

With respect to exon-intron structures, PacBio-ONT inconsistencies mostly come from splice site shifts and skipped short exons due to alignment errors. Such errors appear more often in ONT, although PacBio may also miss exons shorter than 15bp. These inconsistencies increase as intron chains become longer.

The common eukaryotic GTNNGT donor motif is the most frequent cause of donor shifts. However, the probability of incorrectly detecting a donor site in spliced alignments depends on the proximity of GT dinucleotide, rather than motif frequency in the genome. For acceptor sites, this dependency becomes less visible and the overall inconsistency rate is less than for donors. 3bp shifts at NAGNAG acceptors are often seen in both technologies suggesting that unlike other shifts they are not caused by misalignments.

As using only barcoded ONT reads creates a bias towards high-quality reads, we also analyzed all ONT reads. Low-quality reads are shorter, cover fewer introns, and disagree with the annotation more frequently than reads with high Phred scores. Other trends detected in RT event pairs comparison, such as higher inconsistency for long intron chains and intron shifts in the proximity of canonical dinucleotides, are generally preserved and therefore useful to non-barcoded approaches.

We implemented the above observations in a tool for correcting individual read alignments based on gene annotation. Using simulated Nanopore reads we demonstrate that correcting splice site coordinates and misaligned microexons with our tool has a noticeable positive effect on subsequent transcript detection using StringTie2. Moreover, annotation-based correction improves discovery of novel transcripts as they often share exons with known isoforms. Thus, the described findings can be of use for other researchers developing novel algorithms for long-read transcriptome analysis.

Overall, the single-reverse-transcription event approach provides a powerful instrument for platform comparisons. In contrast to the comparisons of distinct molecules, this method offers tertium-non-datur reasoning, where disagreements are known to be caused by errors of one of the platforms.

## Methods

### Experimental details

No new samples or sequencing data was generated for this study, however, we provide a brief description of the samples used and experimental protocols followed in the Supplementary Material. The description is taken from Joglekar et al., 2021. Read statistics are shown in Supplementary Table 1 and 2.

### Alignment to the reference genome using minimap2

PacBio reads were mapped to GRCm38 mouse reference genome with minimap2 v2.17 (Li 2018) using -t 16 -a -x splice:hq --secondary=no options. ONT reads were mapped with -t 16 -a -x splice -k 14 --secondary=no options. When Gencode M21 mouse annotation was used for read mapping, it was converted to BED format using paftools.js gff2bed command (included in minimap2 package) and provided to minimap2 using --junc-bed option.

### Alignment to the genome using deSALT

PacBio reads were mapped to GRCm38 mouse reference genome with deSALT v1.5.6 (Liu et al. 2019) using -t 16 -x ccs options. ONT reads were mapped with -t 16 -x ont1d options.

### Alignment to the genome using GraphMap2

Reads were mapped to GRCm38 mouse genome with GraphMap2 v0.6.5 (Marić et al. 2019) using -t 16 -x rnaseq options. In addition, GraphMap2 was run with Gencode M21 mouse annotation provided using --gtf option, however, the run failed with an error.

### Pairwise read alignment

Pairwise read alignment was performed using the Smith-Waterman local alignment algorithm implemented in SSW python library (Zhao et al. 2013) with default options.

### Sequencing error rate

Sequencing error rates were computed based on minimap2’ alignments using a 3-way comparison between the reference genome and RT read pairs. An error at a certain position in a read from RT pair was reported only when the alignment shows a difference from the genome (i.e. insertion, deletion, or substitution), while the second read from the pair either matches the genome or contains an alternative discrepancy at this position (e.g., another base is inserted). Identical differences from the reference genome (same position and nucleotide) detected in both PacBio and ONT reads from an RT pair were not classified as sequencing errors. An error is deemed to occur within a homopolymer region if any 3bp window in the genome that contains an error position consists of the same nucleotides.

### K-mer identity and homopolymer compression

K-mer identity (k=14bp) with the reference genome was calculated using minimap2 alignments for each exon individually. We first extracted all genomic k-mers from the respective mapped region (of the exon) and then calculated the fraction of the k-mers that occurred within this exon in the read. Homopolymer compression (Au et al. 2012) was performed by substituting all stretches of identical nucleotides (2bp and longer) with a single nucleotide of the same kind in both read sequence and reference sequence from the respective mapping region. K-mer identity was then computed in the same way as for non-compressed sequences.

### TSS / polyA analysis

TSS and polyA sites were assigned to each read as previously done (Joglekar et al. 2021). Specifically, we assigned the closest previously published TSS (Lizio et al. 2015) within 100 bp of the 5’ end of the read mapping. Likewise, we assigned the closest published polyA site (Herrmann et al. 2019) within 100 bp of the 3’ end of the read mapping.

### Intron chain comparison and inconsistency detection

Intron chains were compared against each other as ordered lists of coordinate pairs. In the precise intron chain comparison, two introns are considered equal if their splice site coordinates are identical (delta=0bp), while in the inexact comparison each splice site is allowed to differ by delta=6bp at most. Intron chains are considered as agreeing if they are equal or one chain is a sub-chain of another with respect to the given delta value and disagreeing otherwise.

To detect inconsistencies between reads in RT pairs, intron chains for both reads were extracted from the BAM files obtained with minimap2 and compared against each other as described above. Similarly, to detect agreement between a read and the annotation, a read intron chain extracted from the BAM file was compared against intron chains of known transcripts from Gencode M21 comprehensive mouse annotation. Read is deemed to be consistent if its intron chain agrees with at least one annotated transcript and inconsistent otherwise. Reads that do not overlap with annotated exons (i.e. entirely map to intergenic or intronic regions) are considered as non-informative and are ignored in the analysis.

### Classification of splicing modifications

To classify splice site inconsistencies with the gene annotation reads were assigned to known transcripts using a custom script assign_reads.py available at https://github.com/ablab/platform_comparison, which assigns mapped reads to known isoforms based on intron chains and nucleotide identity. For PacBio reads the script was run with --data_type pacbio_ccs option, for ONT reads --data_type nanopore was used. For further analysis, we selected unambiguous assignments with respective reported splicing modifications (skipped exons, alternative donor or acceptor site). To investigate alternative donors and acceptors (i.e. shift frequency, nucleotide content) only introns with canonical splice sites were used (GT-AG).

The output of the script was also used to track the origin of inconsistent non-barcoded reads. To detect reads having at least one intron entirely located within an annotated exon we selected uniquely assigned reads having this specific type of inconsistency (additional novel intron according to our categories).

### Splice site analysis for non-barcoded reads

To analyze splice site consistency in non-barcoded reads we assigned each read intron separately to an annotated intron (rather than the entire intron chain) with a loose threshold delta=10bp. Such an approach allows to maximize the number of investigated splice sites and consider individual introns even from inconsistent chains. For the analysis, we selected only cases when both read and annotation intron have canonical splice sites (GT-AG). We say that an assigned read intron correctly detects a splice site if its position is equal to the annotated splice site (0bp difference), and incorrectly otherwise. The inconsistency rate of an annotated splice site is defined as the number of incorrect calls divided by the total number of read introns assigned. Each annotated splice site was classified according to the distance to the nearest GT (for donors) or AG (for acceptors) in the vicinity of 10bp. It allowed us to compute overall inconsistency rates for splice sites with respect to this distance.

### Splice site correction algorithm

The correction algorithm takes aligned reads and genome annotation as input. Each read is processed individually, as opposed to the classic transcript construction method that relies on clustering and splice site consensus (Kovaka et al. 2019; Wyman et al. 2020; Tang et al. 2020; Sahlin and Medvedev 2021). An aligned read is first assigned to a reference isoform based on inexact intron chain matching and exon similarity as described above. Further, each read is examined with respect to the accuracy of the detected intron structure. Coordinates of corrected alignments are output in BED12 format. The algorithm is available at https://github.com/ablab/platform_comparison (correct_splice_sites.py).

### Splice site correction algorithm: restoring skipped exons from neighboring splice sites

A reference exon is considered to be skipped during the alignment if it is: (i) shorter than 50bp; (ii) spanned by a read intron; (iii) adjacent exons in the alignment contain extra sequences reaching into the annotated introns surrounding the reference exon and these two extra sequences are of a similar total length as the reference exon (Fig. S8).

### Splice site correction algorithm: correcting individual splice sites

With the same considerations as above, an individual spice site in the read is to be corrected if (1) it is no further than delta=6bp apart from a known splice site and (2) the read alignment has indels close to this position.

### Simulating ONT data

To simulate ONT data we employed the NanoSim software in transcriptome mode (Hafezqorani et al. 2020) using the pre-trained ONT cDNA error model. However, examining the code, we found that NanoSim randomly selects a starting position of a read in an mRNA to simulate truncation. This is performed using a uniform distribution, thus assuming that 5’ and 3’ are identical, which is not the case for the real data. To avoid this pitfall, we mapped raw ONT reads to the reference transcripts using minimap2 (with -*x map-ont* option) and estimated the probabilities of the initial sequence being truncated on each side by N% of its length. We thus modified the NanoSim truncation procedure so that reference sequences are clipped according to empirically derived probabilities (Fig. S9). In addition, we turned off the simulation of random decoy reads, which represent the background noise of the sequencing experiment. We simulated 30M ONT reads using transcripts from the Mouse Gencode v26 basic annotation (Frankish et al. 2021). In addition, a 30bp polyA tail was attached to every transcript prior to simulation. Each transcript with at least one generated read was considered as “expressed” and then represented the ground truth.

### Evaluating transcript model construction

We first generated a reduced genome annotation by removing 20% of expressed spliced transcripts. Removed transcripts are considered novel, while expressed transcripts kept in the annotation represent the set of known models. Using these two sets allows to independently evaluate the ability of the algorithm to report known and discover novel isoforms. StringTie2 (Kovaka et al. 2019) results were similarly split into novel and known models based on the information provided in the GTF file and gffcompare (Pertea and Pertea 2020) was further launched to estimate precision and recall.

## Data access

### Data availability

All data used for this study is publicly available on Gene Expression Omnibus (GEO) database under the accession tokens GSE158450 and GSE173748. All data supporting the findings of this study are provided within the paper and its supplementary information. All additional information will be made available upon reasonable request to the authors.

### Code availability

Barcode detection tool is available as GetBarcodes function in the scicorseqr R-package (github.com/noush-joglekar/scisorseqr). All scripts used for data analysis and spliced alignment correction are available at github.com/ablab/platform_comparison.

## Acknowledgments

We thank Alyona Sidorova and Alexandra Bazarova for their aid with the analysis. This work was supported by the Brain Initiative grant (1RF1MH121267-01 to H.U.T), NIGMS (grant 1R01GM135247-01 to H.U.T), St. Petersburg State University, Russia (grant ID PURE 73023672 to A.M. and A.D.P.). Scientific research was performed at the Research park of St.Petersburg State University «Computing Center».

