## Supplementary Material for "Sequencing of individual barcoded cDNAs on Pacific Biosciences and Oxford Nanopore reveals platform-specific error patterns"

### Supplementary Note "Experimental details"

Here we provide a brief description of the samples used and experimental protocols followed (taken from Joglekar et al., 2021). C57BL/6NTac (n=3) female pups were used in generating the data. For the single-cell experiments (P7; n=1), the brains were removed and placed on a stainless-steel brain matrix for mouse (coronal repeatable sections, 1mm spacing), and the hippocampus was dissected in cold PBS solution. After dissection, tissues were snap frozen in dry ice until processing. For 10X Visium experiments (P8; n=1) brains were fresh-frozen and embedded in OCT. Mice were housed in an air-conditioned room at 65-75°F (~18-23°C) with 40-60% humidity and a 12-h-light, 12-h-dark cycle. Animal experiments were approved by the institutional animal care and use committee (IACUC) of Weill Cornell Medicine

**Tissue dissociation and 10x Genomics single-cell capture.**

The dissected hippocampal tissue was dissociated into a single-cell suspension according to the manufacturer's protocol and the disassociated cells were captured on the 10x Genomics Chromium controller according to the Chromium Single Cell 3′ Reagent Kits V2 User Guide with the following modification. PCR cycles were increased, from the recommended ten cycles for recovery of 8,000 cells to 16 cycles to target a yield of cDNA enabling simultaneous Illumina and PacBio library preparation.

**Illumina and Pacific Biosystems library preparation.**

Illumina library preparation was performed using 100 ng of amplified cDNA following the Chromium Single Cell 3′ Reagent Kits V2 User Guide reducing final indexing PCR cycles to ten cycles from the recommended 14 cycles to increase library complexity. Sequencing was performed on HiSeq4000 according to 10x Genomics run mode. PacBio library preparation was performed with 500 ng of amplified cDNA using SMRTbell Express Template Prep Kit V2.0 to obtain Sequel II compatible library complex and was sequenced on a total 16 Sequel I SMRTcells with a run time of 10 hours and 10 Sequel II SMRTcells with a run time of 30 hours across samples and replicates.

**PromethION library preparation and sequencing of cDNA.**

Oxford Nanopore compatible library was produced using 350ng of either Sage Blue Pippin size-selected cDNA or non-selected cDNA derived from 10x Genomics following the Genomic DNA by Ligation protocol from Oxford Nanopore. Loading inputs on the PromethION was increased to 150fmol and sequenced for 20 hours, and base-calling was done using Guppy (3.2.10).

**Generation of circular consensus reads.**

Using the default SMRT-Link parameters, we performed circular consensus sequencing (CCS) with IsoSeq3 with the following modified parameters: maximum subread length 14,000bp, minimum subread length 10bp, and minimum number of passes 3.

**Barcode detection.**

Post filtering for quality control, a list of 16mer barcodes corresponding to single cells or spots was obtained from 10X sequencing output. Assuming an error rate of 10%, PolyA tails were identified by obtaining the position of 9 consecutive A's or T's in the last 100nt of PacBio CCS and ONT reads respectively. Using the position of the PolyA tail, a sliding window was used to identify perfect matching barcodes in the 36 bp window preceding the PolyA tail, and UMIs were identified as the 10 or 12 bp (depending on the dataset) following the barcode. This method is implemented in the scisorseqr package under the GetBarcodes function. Read statistics are shown in Supplementary Tables 1 and 2.

| | PacBio | ONT |
|---|---|---|
| **# reads** | 3,371,331 | 73,181,790 |
| **# spliced reads** | 1,266,498 | 28,961,005 |
| **# reads with poly-A tails** | 3,321,897 | 38,986,771 |
| **# of unique barcodes detected** | 3,024 | 3,023 |
| **# reads with detected barcodes** | 2,873,455 | 12,153,599 |
| **# common molecules** | 54,752 | |

**Supplementary Table 1. Read statistics on the entire Sl-ISO-Seq dataset.** For barcode detection, we used a list of 3,024 16mer barcodes obtained from 283,600,728 10X reads.

| | PacBio | ONT |
|---|---|---|
| **# reads** | 38,290,265 | 29,212,434 |
| **# spliced reads** | 18,223,425 | 9,799,668 |
| **# reads with poly-A tails** | 23,020,473 | 7,830,696 |
| **# of unique barcodes detected** | 6,921 | 6,921 |
| **# reads with detected barcodes** | 15,969,422 | 1,741,924 |
| **# common molecules** | 274,287 | |

**Supplementary Table 2. Read statistics on the entire sc-ISOr-Seq dataset.** For barcode detection, we used a list of 6,922 16mer barcodes obtained from 357,009,685 10X reads.

| | minimap2 | GraphMap2 | deSALT |
|---|---|---|---|
| **Total running time (sec)** | 433 | 7331 | 500 |
| **# RT pairs** | 54,752 | 52,927 | 57,040 |
| **Median alignment length (PacBio)** | 888 | 870 | 892 |
| **Median alignment length (ONT)** | 877 | 856 | 882 |
| **Median difference (PacBio read is longer)** | 14 | 15 | 14 |
| **Median difference (ONT read is longer)** | 8 | 9 | 8 |

**Supplementary Table 3. Different aligners statistics on the Sl-ISO-Seq dataset.** An RT pair is a pair where ONT and PacBio read have the same barcode and UMI and align to the same gene. The median difference between aligned lengths of PacBio read and ONT read from the RT read pair was calculated separately for cases when PacBio (ONT) alignment was longer.

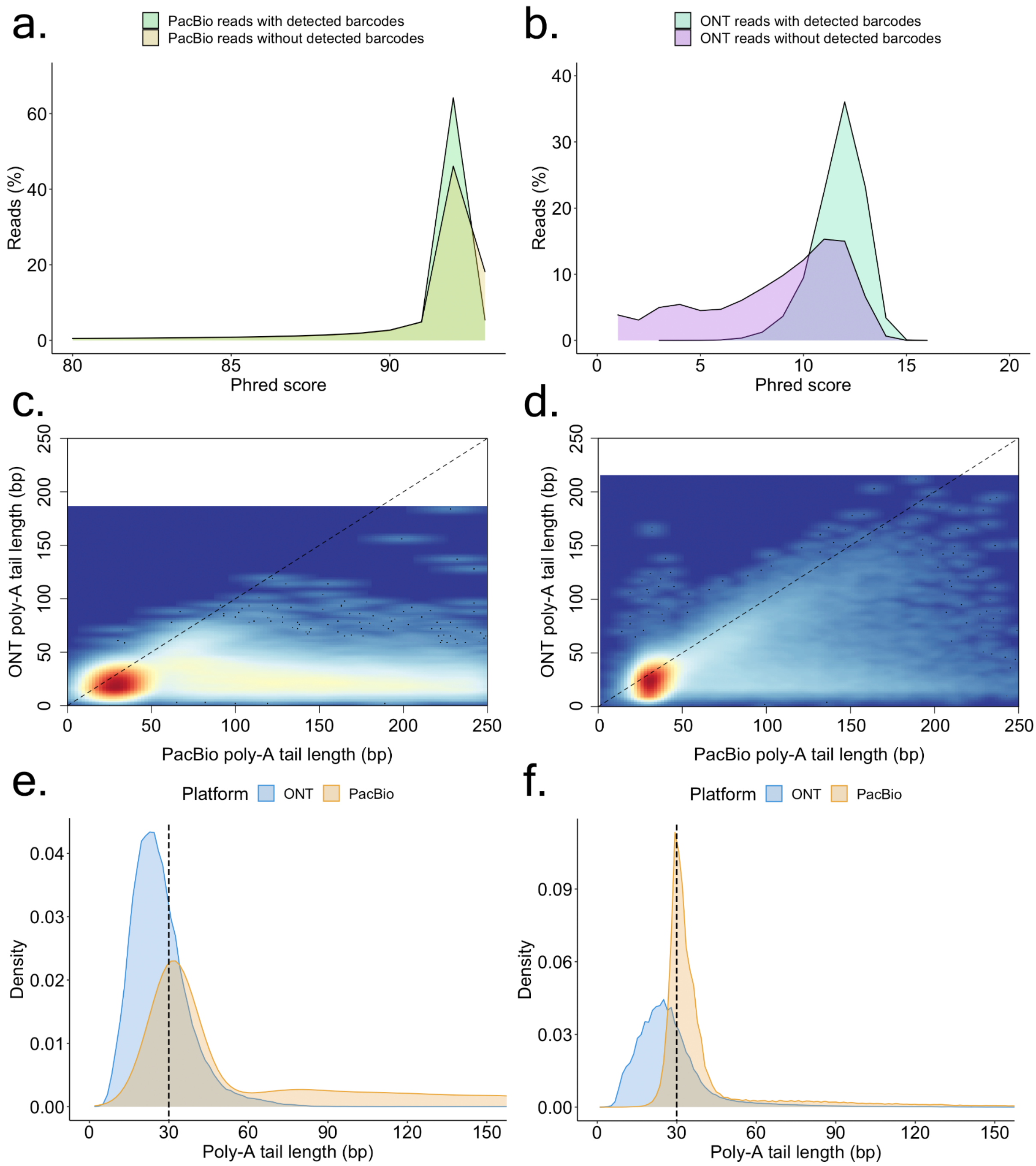


**Fig. S1. Primary read characteristics of ScISOr-Seq and Sl-ISO-Seq data. a.** Phred score distribution for Oxford Nanopore reads from ScISOr-Seq dataset with (light blue) and without (purple) detected barcodes. **b.** Phred score distribution for PacBio CCS reads from ScISOr-Seq dataset with (light green) and without (yellow) detected barcodes. **c.** Heatscatter plot showing the distribution of RT pairs according to the length of a poly-A tail detected in the respective PacBio read (X axis) and ONT read (Y axis) for Sl-ISO-Seq dataset. **d.** Heatscatter plot showing the distribution of RT pairs according to the length of a poly-A tail detected in the respective PacBio read (X axis) and ONT read (Y axis) for ScISOr-Seq dataset. **e.** Density distribution of poly-A tail lengths in PacBio (yellow) and ONT (blue) reads for Sl-ISO-Seq dataset. The dashed line represents the expected length of poly-A tails (30 bp). **f.** Density distribution of poly-A tail lengths in PacBio (yellow) and ONT (blue) reads for ScISOr-Seq dataset. The dashed line represents the expected length of poly-A tails (30 bp).

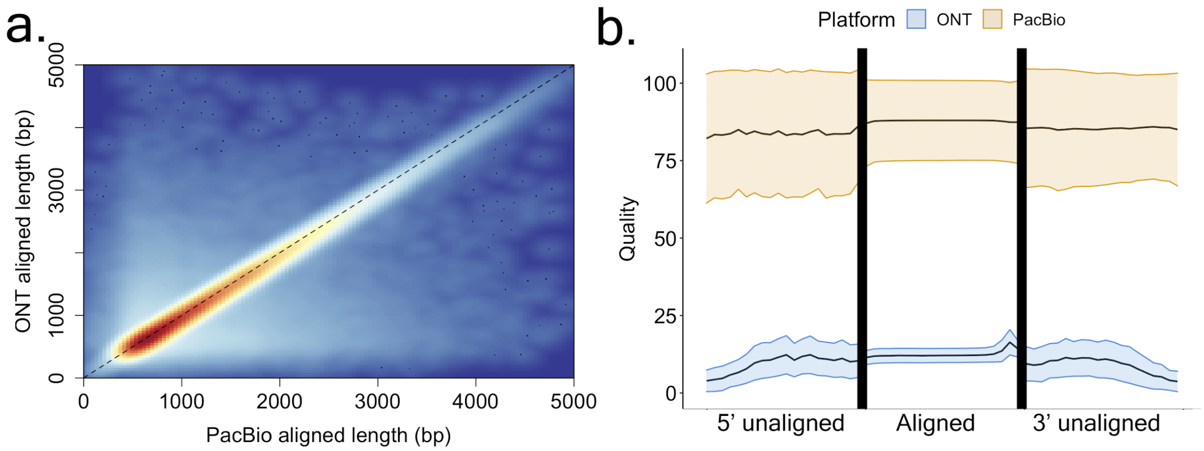


**Figure S2. Alignment characteristics of RT read pairs of ScISOr-Seq data. a.** Heatscatter plot showing aligned lengths of respective PacBio read (X axis) and ONT read (Y axis) from the RT read pair after mapping to the genome using minimap2 for ScISOr-Seq dataset (Spearman Rho: 0.90, $p<2.2*10^{-16}$). **b.** Mean phred score distribution along the reads for aligned (middle) and unaligned (left and right) parts of PacBio (yellow) and ONT (blue) reads based on a (reference-free) pairwise Smith-Waterman alignment of the PacBio and Nanopore reads from the RT read pair for ScISOr-Seq dataset. Lower and upper bounds represent the standard deviation of the Phred score distribution.

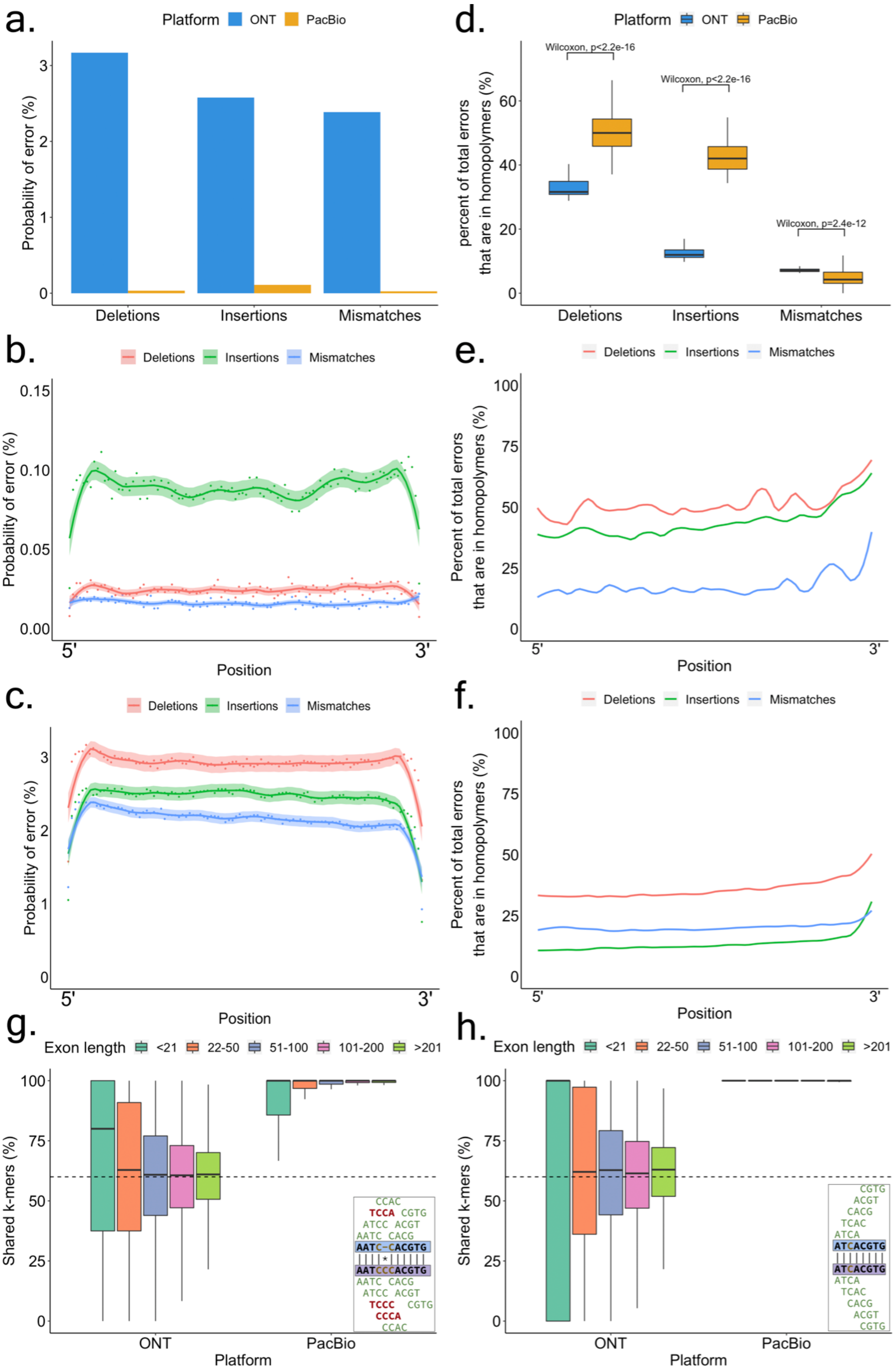
a.
Platform ONT PacBio
Probability of error (%)
Deletions
Insertions
Mismatches
d.
Platform ONT PacBio
Wilcoxon, p<2.2e-16
Wilcoxon, p<2.2e-16
Wilcoxon, p=2.4e-12
percent of total errors that are in homopolymers (%)
Deletions
Insertions
Mismatches
b.
Deletions Insertions Mismatches
Probability of error (%)
5'
3'
Position
e.
Deletions Insertions Mismatches
Percent of total errors that are in homopolymers (%)
5'
3'
Position
c.
Deletions Insertions Mismatches
Probability of error (%)
5'
3'
Position
f.
Deletions Insertions Mismatches
Percent of total errors that are in homopolymers (%)
5'
3'
Position
g.
Exon length <21 22-50 51-100 101-200 >201
Shared k-mers (%)
ONT
PacBio
Platform
h.
Exon length <21 22-50 51-100 101-200 >201
Shared k-mers (%)
ONT
PacBio
Platform

**Fig. S3. Sequencing error rates and k-mer identity for PacBio and Nanopore from SI-ISO-Seq dataset. a.** Frequency of each sequencing error type in PacBio (yellow) and ONT reads (blue). **b.** Sequencing error probability with respect to the position in read for PacBio data. **c.** Sequencing error probability with respect to the position in read for ONT data. **d.** Percent of sequencing errors located in homopolymers for each error type in PacBio (yellow) and ONT reads (blue). **e.** Fraction of sequencing errors located in homopolymers with respect to position in the read for PacBio data. **f.** Fraction of sequencing errors located in homopolymers with respect to position in the read for ONT data. **g.** K-mer identity (k=14) of separate exons for ONT (left) and PacBio reads (right) with respect to the reference exon length. *Bottom right:* An example of two sequences and their shared (green) and distinct *k*-mers (red) before homopolymer compression. **h.** K-mer identity (k=14) of separate exons after homopolymer compression for ONT (left) and PacBio reads (right) with respect to the reference exon length. *Bottom right:* An example of the same two sequences from Fig. 3g and their shared (green) and distinct *k*-mers (red) after homopolymer compression. phred score distribution.

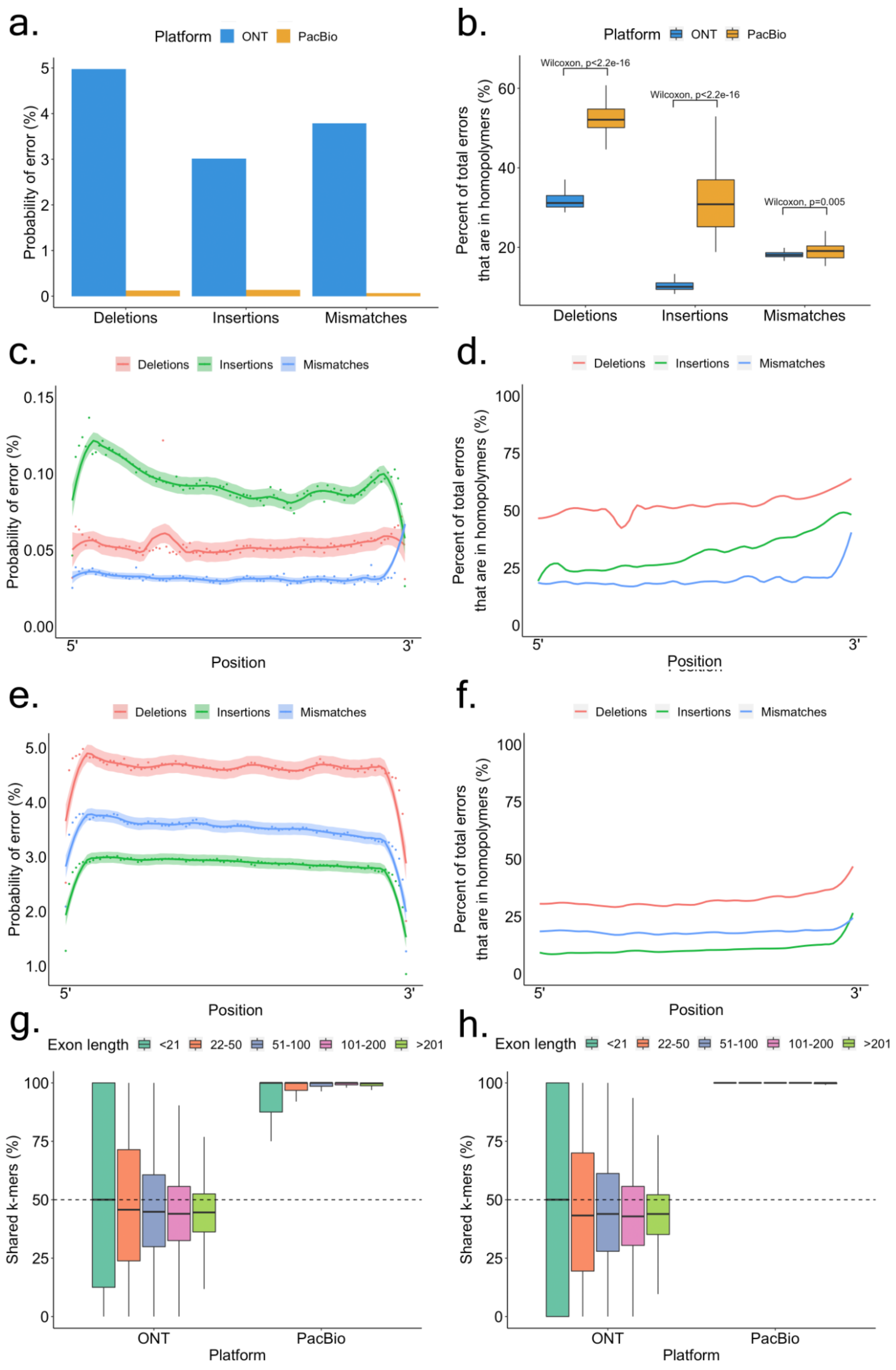
a.
Platform ONT PacBio
Probability of error (%)
Deletions
Insertions
Mismatches
b.
Platform ONT PacBio
Percent of total errors that are in homopolymers (%)
Wilcoxon, p<2.2e-16
Wilcoxon, p<2.2e-16
Wilcoxon, p=0.005
Deletions
Insertions
Mismatches
c.
Deletions Insertions Mismatches
Probability of error (%)
5'
3'
Position
d.
Deletions Insertions Mismatches
Percent of total errors that are in homopolymers (%)
5'
3'
Position
e.
Deletions Insertions Mismatches
Probability of error (%)
5'
3'
Position
f.
Deletions Insertions Mismatches
Percent of total errors that are in homopolymers (%)
5'
3'
Position
g.
Exon length <21 22-50 51-100 101-200 >201
Shared k-mers (%)
ONT
PacBio
Platform
h.
Exon length <21 22-50 51-100 101-200 >201
Shared k-mers (%)
ONT
PacBio
Platform

**Fig. S4. Sequencing error rates and k-mer identity for PacBio and Nanopore from ScISOr-Seq dataset. a.** Frequency of each sequencing error type in PacBio (yellow) and ONT reads (blue). **b.** Percent of sequencing errors located in homopolymers for each error type in PacBio (yellow) and ONT reads (blue). **c.** Sequencing error probability with respect to the position in read for PacBio data. **d.** Fraction of sequencing errors located in homopolymers with respect to position in the read for PacBio data. **e.** Sequencing error probability with respect to the position in read for ONT data. **f.** Fraction of sequencing errors located in homopolymers with respect to position in the read for ONT data. **g.** K-mer identity (k=14) of separate exons for ONT (left) and PacBio reads (right) with respect to the reference exon length. **h.** K-mer identity (k=14) of separate exons after homopolymer compression for ONT (left) and PacBio reads (right) with respect to the reference exon length.

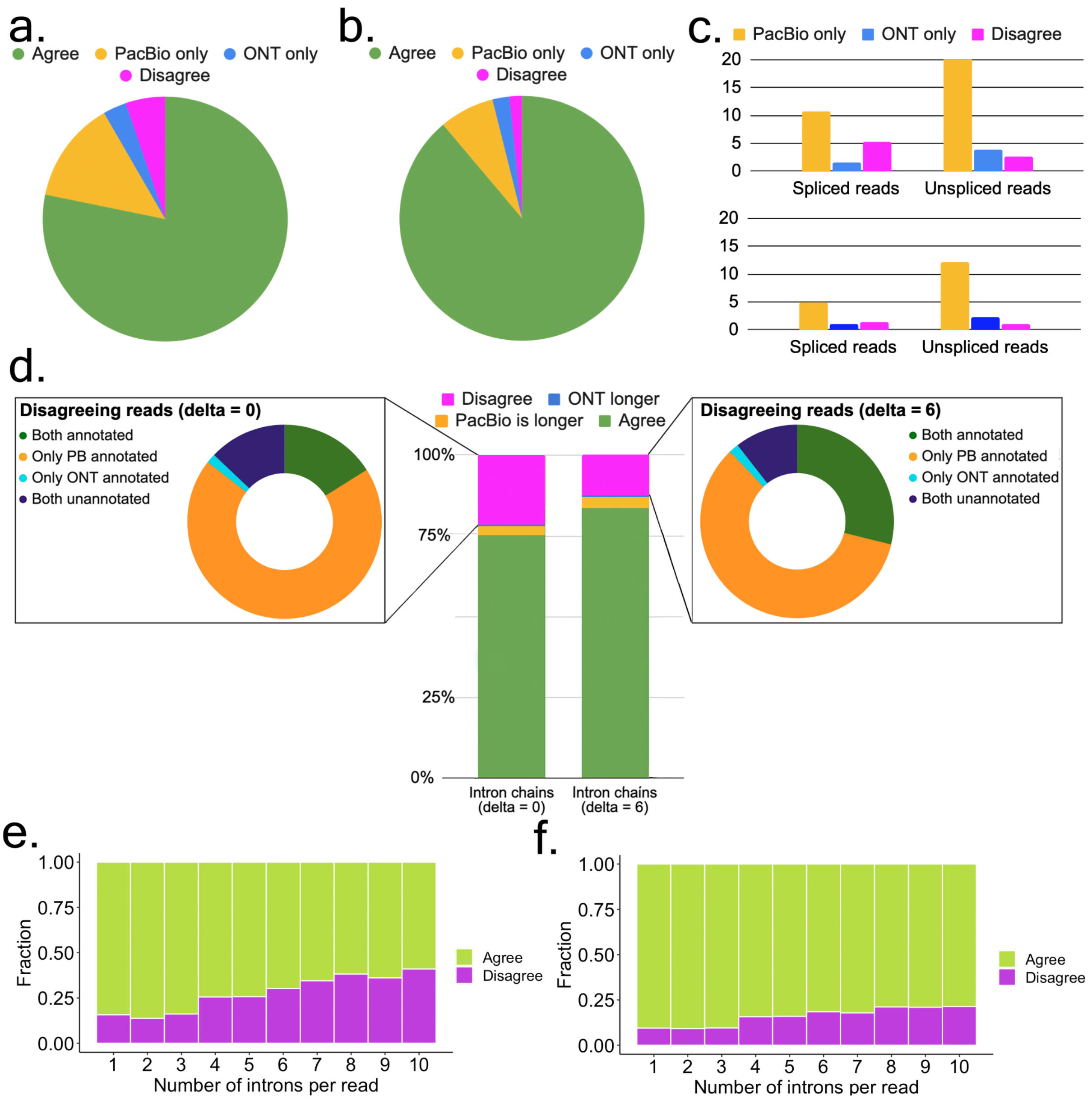


**Figure S5. Agreement in RT read pairs of ScISOr-Seq data. a.** Fractions of TSS assignments that agree (green), disagree (magenta), or found only in one read (blue for ONT, yellow for PacBio) from the RT pair. **b.** Same as Fig. S5a, but for Poly-A sites. **c.** Percentage of RT read pairs that disagree on the assigned TSS (top) and Poly-A site (bottom): only PacBio read was assigned (yellow), only ONT (blue), both assigned but to different sites (magenta). **d.** *Middle*: fraction of intron chains from the RT read pairs that agree (green), disagree (magenta), or one chain being longer (blue for ONT, yellow for PacBio) when splice junctions are compared precisely (left, delta=0bp) or inexactly (right, delta=6bp). *Top left:* classification of disagreeing intron chains from RT read pairs with respect to the reference annotation (delta=0bp): both are inconsistent with the annotation (dark blue), both correspond to known (different) transcripts despite the disagreement (green), PacBio is consistent with the annotation while ONT chain is not (yellow) and vise versa (light blue). *Top right:* classification of disagreeing intron chains from RT read pairs with respect to the reference annotation using inexact comparison (delta=6bp). **e.** Fraction of agreeing (green) and disagreeing (magenta) intron chains with respect to intron chain length when compared precisely (delta=0bp). **f.** Fraction of agreeing (green) and disagreeing (magenta) intron chains with respect to intron chain length when compared with delta=6bp.

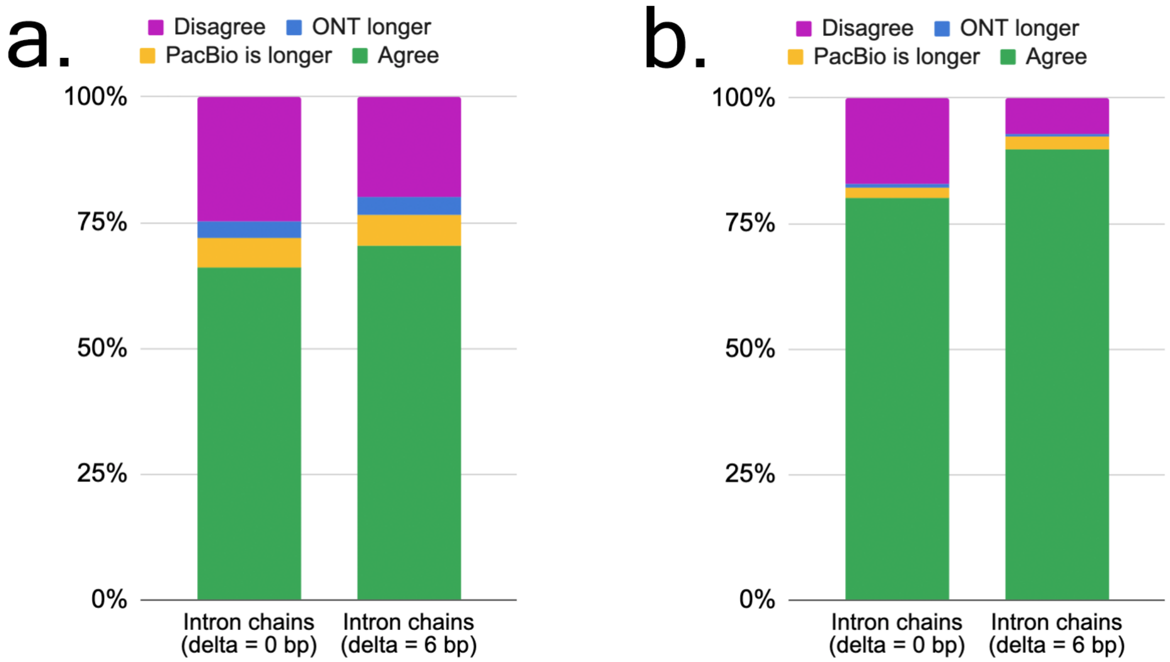


**Figure S6. Agreement in RT read pairs of Sl-ISO-Seq data using alignments produced by alternative aligners.** The corresponding plot for minimap2 alignments is shown in Fig. 3d, in 81.8% (89%) of RT read pairs the PacBio and the ONT read had identical intron chains with delta=0bp (6bp). **a.** Fraction of intron chains from the RT read pairs that agree (green), disagree (magenta), or one chain being longer (blue for ONT, yellow for PacBio) when splice junctions are compared precisely (left, delta=0bp) or inexactly (right, delta=6bp). Read alignments were produced by GraphMap2. In 66.1% (69.2%) of RT read pairs the PacBio and the ONT read had identical intron chains with delta=0bp (6bp). **b.** Same as Fig. 5a, but for alignments produced by deSALT. In 79.8% (89.8%) of RT read pairs the PacBio and the ONT read had identical intron chains with delta=0bp (6bp).

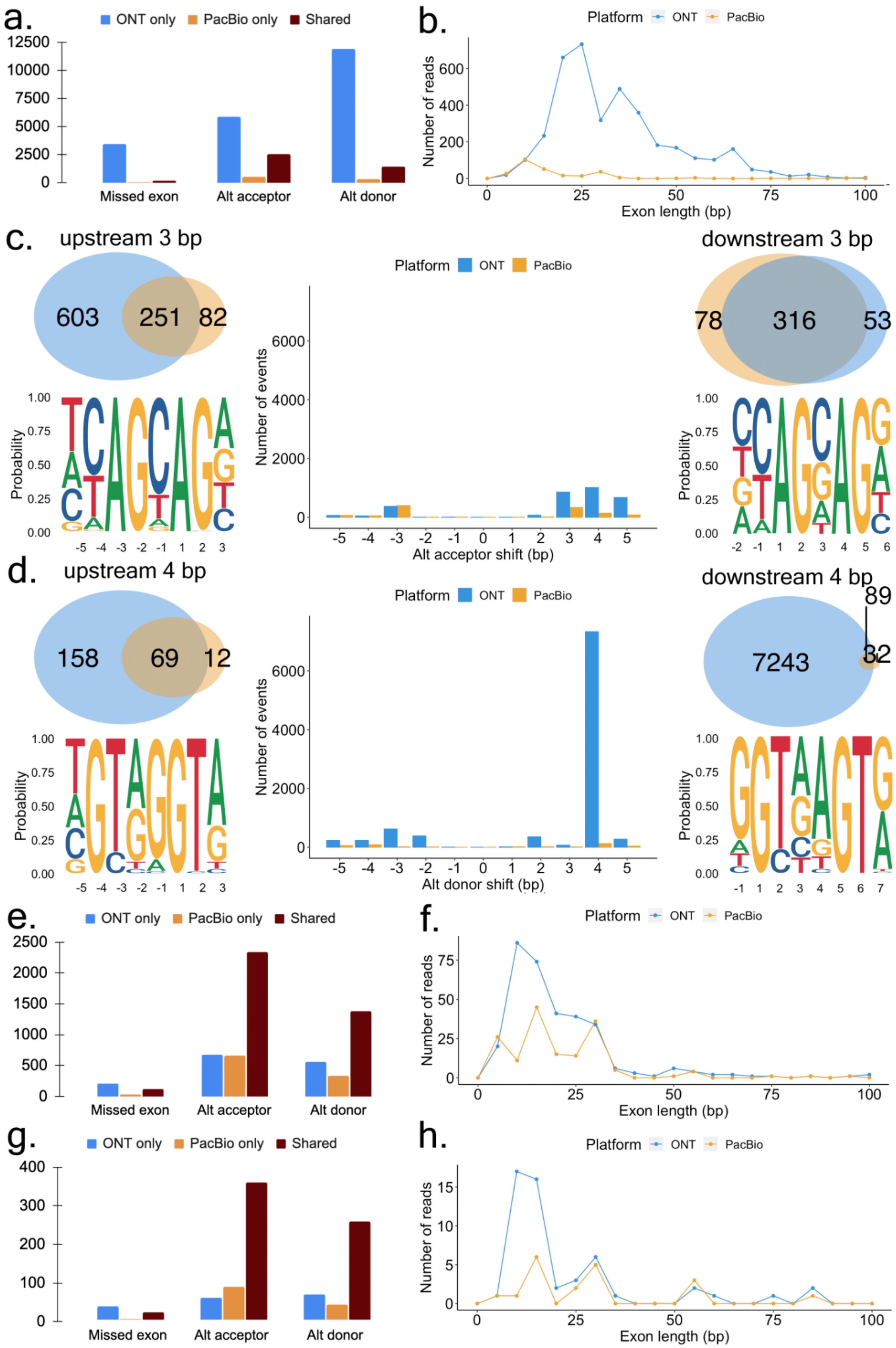

a.
ONT only
PacBio only
Shared
Missed exon
Alt acceptor
Alt donor
b.
Platform
ONT
PacBio
Number of reads
Exon length (bp)
c.
upstream 3 bp
603
251
82
downstream 3 bp
78
316
53
Probability
Number of events
Alt acceptor shift (bp)
d.
upstream 4 bp
158
69
12
downstream 4 bp
89
7243
32
Alt donor shift (bp)
e.
f.
g.
h.

**Figure S7. Exon and splice site characteristics underlying disagreements between PacBio and Nanopore in ScISOr-Seq data (a-f) and Sl-ISO-Seq data (g-h). a.** Number of missed exons (left), alternative acceptors (middle), and donors (right) with respect to the reference annotation that occur only in ONT read (blue), only in PacBio read (yellow), and in both reads from RT pair (brown). **b.** Length distribution for skipped exons in PacBio reads (yellow) and ONT reads (blue). **c.** *Middle:* number of alternative acceptor sites in PacBio reads (yellow) and ONT reads (blue) with respect to distance from the annotated acceptor site. *Top left:* Venn diagram for 3bp upstream alternative acceptor sites in PacBio (yellow) and ONT reads (blue) from RT pair. *Bottom left:* Nucleotide frequency for loci where 3bp upstream acceptor sites occur. *Top right:* Venn diagram for 3bp downstream alternative acceptor sites in PacBio (yellow) and ONT reads (blue) from RT pair. *Bottom right:* Nucleotide frequency for loci where 3bp downstream acceptor sites occur. **d.** *Middle:* number of alternative donor sites in PacBio reads (yellow) and ONT reads (blue) with respect to distance from the annotated donor site. *Top left:* Venn diagram for 4bp upstream alternative donor sites in PacBio (yellow) and ONT reads (blue) from RT pair. *Bottom left:* Nucleotide frequency for loci where 4bp upstream donor sites occur. *Top right:* Venn diagram for 4bp downstream alternative donor sites in PacBio (yellow) and ONT reads (blue) from RT pair. *Bottom right:* Nucleotide frequency for loci where 4bp downstream donor sites occur. **e.** Number of missed exons (left), alternative acceptors (middle), and donors (right) with respect to the reference annotation that occur only in ONT read (blue), only in PacBio read (yellow), and in both reads from RT pair (brown) in ScISOr-Seq data, the genome annotation was used during the alignment step. **f.** Length distribution for skipped exons in PacBio reads (yellow) and ONT reads (blue) in ScISOr-Seq data, the genome annotation was used during the alignment step. **g.** Number of missed exons (left), alternative acceptors (middle), and donors (right) with respect to the reference annotation that occur only in ONT read (blue), only in PacBio read (yellow), and in both reads from RT pair (brown) in Sl-ISO-Seq data, the genome annotation was used during the alignment step. **h.** Length distribution for skipped exons in PacBio reads (yellow) and ONT reads (blue) in Sl-ISO-Seq data, the genome annotation was used during the alignment step.

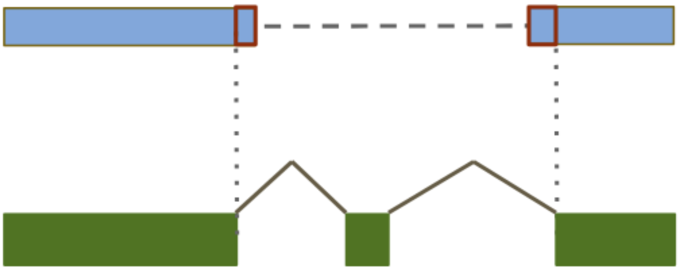

**Figure S8. Misaligned short exon.** An ONT read alignment (blue) misses a short reference exon (green) by extending an alignment of the neighboring exons (red).

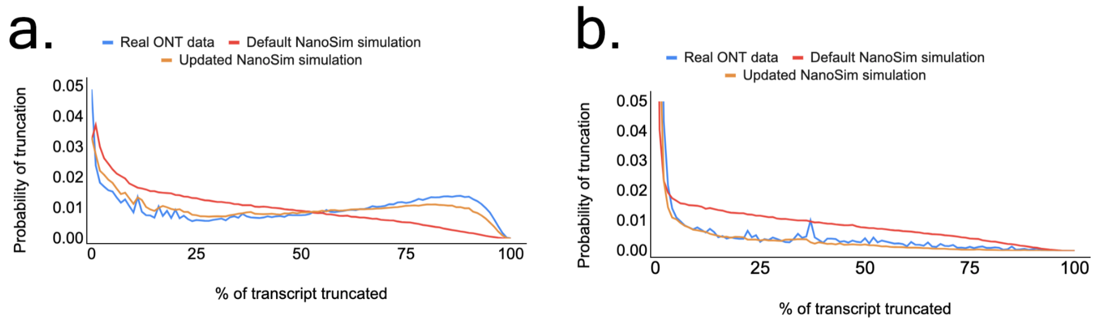


**Figure S9. Truncation probabilities for real and simulated ONT data.** (a) Probability of an ONT read being truncated by X% on the 5’ end for real Sl-ISO-seq data (red), simulated using NanoSim with default settings (blue), and simulated using NanoSim with improved truncation procedure (green). Truncated fractions were estimated by mapping reads onto reference transcriptome using minimap2 with -x map-ont option. (b) Same as S9a but for the 3’ end.
